# Hepatoprotective effects of Aureobasidium pullulans derived Beta 1,3-1,6 biological response modifier glucans in a STAM- animal model of non-alcoholic steatohepatitis

**DOI:** 10.1101/2021.07.08.451700

**Authors:** Nobunao Ikewaki, Gene Kurosawa, Masaru Iwasaki, Senthilkumar Preethy, Vidyasagar Devaprasad Dedeepiya, Suryaprakash Vaddi, Rajappa Senthilkumar, Gary A Levy, Samuel JK Abraham

## Abstract

**Background:** Non-alcoholic fatty liver disease (NAFLD) and non-alcoholic steatohepatitis (NASH) are highly prevalent conditions characterized by inflammation and fibrosis of the liver which can progress to cirrhosis and hepatocellular carcinoma if left untreated. Lifestyle disorders such as obesity, diabetes and dyslipidaemia predispose to and are associated with the disease progression. Conventional modalities are mainly symptomatic, with no definite solution. Beta glucan-based biological response modifiers are a potential strategy in lieu of their beneficial metabolic effects. *Aureobasidium pullulans* strains AFO-202 and N-163 beta glucans were evaluated for anti-fibrotic and anti-inflammatory hepatoprotective potentials in a NASH animal model in this study.

**Methods:** In the STAM™ murine model of NASH, five groups were studied for eight weeks— (1) vehicle (RO water), (2) AFO-202 beta glucan; (3) N-163 beta glucan, (4) AFO-202+N-163 beta glucan, and (5) telmisartan (standard pharmacological intervention). Evaluation of biochemical parameters in plasma and hepatic histology including Sirius red staining and F4/80 immunostaining were performed.

**Results:** AFO-202 beta glucan significantly decreased inflammation-associated hepatic cell ballooning and steatosis. N-163 beta glucan decreased fibrosis and inflammation significantly (p value<0.05). The combination of AFO-202 with N-163 significantly decreased the NAFLD Activity Score (NAS) compared with other groups.

**Conclusion:** This preclinical study supports the potential of N-163 and AFO-202 beta glucans alone or in combination as potential preventive and therapeutic agent(s), for NASH.

**Graphical abstract:** 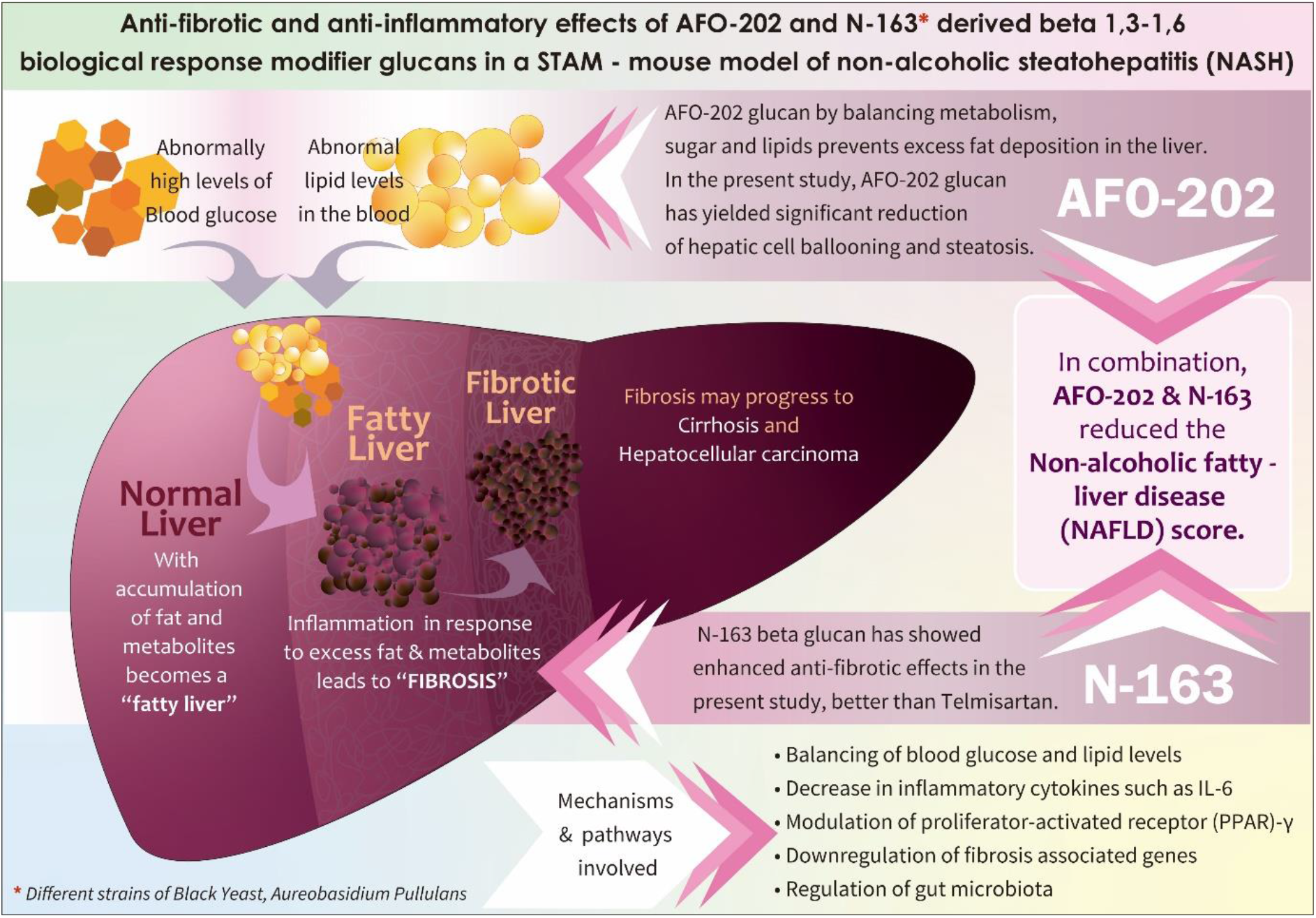

## Introduction

Non-alcoholic fatty liver disease (NAFLD) refers to a group of conditions in which there is excess fat accumulation on the liver in people who drink little or no alcohol [1]. Non-alcoholic steatohepatitis (NASH) is a severe form of NAFLD. NAFLD or NASH progresses to liver fibrosis, liver cirrhosis, liver failure or carcinoma if not treated. Increased prevalence of obesity and metabolic syndrome, diabetes and dysregulated lipid levels, all add to the problem of NAFLD and NASH. NAFLD and NASH involve pathologic features such as hepatic steatosis, lobular inflammation, hepatocellular ballooning and liver fibrosis, which ultimately lead to cirrhosis [1,2]. There are no definite treatments for NASH. Conventional approaches aim to address the underlying condition such as diabetes and metabolic disease with lifestyle changes, weight reduction, specific medication such as thiazolidinediones, lipid-lowering agents, cytoprotective agents and antioxidants such as vitamin E [3]. Angiotensin receptor blockers (ARBs) such as telmisartan, which act by modulating transcription factor peroxisome proliferator-activated receptor (PPAR)-γ activity [4], thereby increasing insulin sensitivity, are increasingly being advocated. However, the underlying aetiology and disease pathogenesis need more holistic approaches.

Animal models have proven to be highly useful to investigate the etiopathogenesis of a number of human diseases [5]. Stelic Animal Model (STAM™) is an animal model that recapitulates the disease progression of that which occurs in human NASH/HCC [6]. In this model, C57BL/6 mice aged two days are given a single dose of streptozotocin to reduce the insulin secretory capacity. When the mice turn four weeks of age they are started on a high-fat diet feeding. This model has a background of late type 2 diabetes which progresses into fatty liver, NASH, fibrosis and consequently HCC [2,6]. In this study we have employed the STAM – animal model to study the hepatoprotective anti-fibrotic and anti-inflammatory effects of beta glucans from a black yeast, *Aureobasidium pullulans*. Beta glucans are potent biological response modifiers that have been proven to be effective in modulating dysregulated metabolism by regulating blood glucose and lipid levels. The *Aureobasidium pullulans* AFO-202 strain-derived 1,3-1,6 beta glucan has been demonstrated to decrease HbA1c to normal values and decrease fasting and post-prandial blood glucose in human clinical studies [7,8]. This GMP-manufactured beta glucan has been proven to regulate lipid levels of triglycerides, total cholesterol and HDL cholesterol in another human clinical study [9]. Another variant of the 1,3-1,6 beta glucan has been derived from a novel strain, N-163 of *Aureobasidium pullulans*, which in in vitro studies has shown to have a positive effect on lipid metabolism (data unpublished). In the present study, we report the anti-fibrotic and anti-inflammatory hepatoprotective effects of AFO-202 and N-163-strains derived beta glucan individually and in combination, in STAM mice.

## Materials and methods

### Mice

C57BL/6J mice were obtained from Japan SLC, Inc. (Japan). All animals used in this study were cared for under the following guidelines: Act on Welfare and Management of Animals (Ministry of the Environment, Japan, Act No. 105 of October 1, 1973), standards relating to the care and management of laboratory animals and relief of pain (Notice No.88 of the Ministry of the Environment, Japan, April 28, 2006) and the guidelines for proper conduct of animal experiments (Science Council of Japan, June 1, 2006). Protocol approvals were obtained from SMC Laboratories, Japan’s IACUC (Study reference no: SP_SLMN128-2107-6_1). Mice were maintained in a specific pathogen-free (SPF) facility under controlled conditions of temperature (23 ± 3°C), humidity (50 ± 20%), lighting (12-hour artificial light and dark cycles; light from 8:00 to 20:00) and air exchange.

The STAM model of NASH was induced as previously described [5]. Mice were given a single subcutaneous injection of 200 μg streptozotocin (STZ, Sigma-Aldrich, USA) solution 2 days after birth and fed with a high-fat diet (HFD, 57 kcal% fat, Cat# HFD32, CLEA Japan, Inc., Japan) from 4 – 9 weeks of age [2, 6]. All mice develop liver steatosis and diabetes and at 3 weeks mice had established steatohepatitis, histologically [2].

#### Study groups

There were five study groups, described below. Eight mice were included in each study group.

##### Group 1: Vehicle

Eight NASH mice were orally administered the vehicle (RO water) in a volume of 5 mL/kg once daily from 6 to 9 weeks of age.

##### Nichi Glucan groups

The dose of Nichi Glucan was decided based on the earlier studies of AFO-202 strain derived beta glucan in human healthy volunteers and subjects with lifestyle disorders (diabetes dyslipidaemia) [6-8] and N-163 strain derived beta glucan in healthy volunteers (data unpublished).

##### Group 2: AFO-202 Beta Glucan

Eight NASH mice were orally administered the vehicle supplemented with AFO-202 beta glucan at a dose of 1 mg/kg in a volume of 5 mL/kg once daily from 6 to 9 weeks of age.

##### Group 3: N-163 Beta Glucan

Eight NASH mice were orally administered the vehicle supplemented with N-163 beta glucan at a dose of 1 mg/kg in a volume of 5 mL/kg once daily from 6 to 9 weeks of age.

##### Group 4: AFO-202 Beta Glucan + N-163 Beta Glucan

Eight NASH mice were orally administered the vehicle supplemented with AFO-202 beta glucan at a dose of 1 mg/kg in a volume of 5 mL/kg once daily and orally administered the vehicle supplemented with N-163 beta glucan at a dose of 1 mg/kg in a volume of 5 mL/kg once daily from 6 to 9 weeks of age.

##### Group 5: Telmisartan

Eight NASH mice were orally administered the vehicle supplemented with telmisartan at a dose of 10 mg/kg once daily from 6 to 9 weeks of age.

Telmisartan which has been reported to have antisteatotic, anti-inflammatory and antifibrotic effects in STAM model was used as the positive comparator.

AFO-202 and N-163 beta glucan were provided by GN Corporation Co Ltd. Telmisartan (Micardis^®^) was purchased from Boehringer Ingelheim GmbH (Germany).

### 1.1. Preparation of test substances

#### 1.1.1. AFO-202 Beta Glucan and N-163 Beta Glucan

AFO-202 beta glucan or N-163 beta glucan was mixed in the required amount of RO water and stirred until it completely dissolved. The solution was dispensed into 7 tubes and stored at 4°C until the day of administration. The dosing formulations were stirred prior to administration. The dosing formulations were used within 7 days.

#### 1.1.2. Telmisartan

Formulations were freshly prepared prior to administration. One tablet of telmisartan was transferred into mortar and triturated using a pestle by adding RO water gradually to get 1 mg/mL of homogeneous suspension.

### 1.2. Randomization

NASH model mice were randomized into five groups of eight mice at six weeks of age based on their body weight the day before the start of treatment. The randomization was performed by body weight-stratified random sampling using Microsoft Excel software. NASH model mice were stratified by their body weight to get the SD and difference in the mean weights among groups as small as possible.

### 1.3. Animal monitoring and sacrifice

Mice were monitored for clinical signs (lethargy, twitching, laboured breathing), behaviour and survival. Body weight was recorded daily. Mice were observed for significant clinical signs of toxicity, moribundity and mortality before and after administration. The animals were sacrificed at 9 weeks of age by exsanguination through direct cardiac puncture under isoflurane anaesthesia (Pfizer Inc.). At the time of sacrifice, the mice are expected to have reached the steatohepatitis phase of the disease and a mild hepatic fibrotic stage [2].

If an animal showed >25% body weight loss within a week or >20% body weight loss compared with the previous day, the animal was euthanized ahead of study termination, and samples were not collected. If it showed a moribundity sign, such as prone position, the animal was euthanized ahead of study termination, and samples were not collected.

### 1.4. Sample collection

The following samples were collected and stored.

- Frozen Plasma
- SNAP Frozen liver
- Paraffin-embedded liver
- OCT-embedded liver

#### Preparation of plasma samples

At study termination, non-fasting blood was collected through direct cardiac puncture using pre-cooled syringes. The collected blood was transferred in pre-cooled polypropylene tubes with anticoagulant (Novo-Heparin) and stored on ice until centrifugation. The blood samples were centrifuged at 1,000 x g for 15 minutes at 4°C. The supernatant was collected and stored at −80°C for biochemistry and evaluation.

#### Preparation of liver samples

After sacrifice, the whole liver was collected and washed with cold saline. Photos of individual whole livers (parietal side and visceral side) were taken. Liver weight was measured, and liver-to-body weight ratio was calculated. The left lateral lobes of the livers were separated, dissected and stored.

A. Liver specimens were stored at −80°C embedded in optimal cutting temperature (OCT, Sakura Finetek Japan, Japan) compound for immunohistochemistry.
B. Liver specimens were fixed in Bouin’s solution (Sigma-Aldrich Japan, Japan) for 24 hours. After fixation, these specimens were proceeded to paraffin embedding for HE and Sirius red staining.
C. Liver specimens were snap frozen in liquid nitrogen and stored at −80°C for further analysis.

The left and right medial lobes were snap frozen in liquid nitrogen and stored at −80°C for evaluation.

The right lobe was snap frozen in liquid nitrogen and stored at −80°C for biochemistry analysis.

The caudate lobe was snap frozen in liquid nitrogen and stored at −80°C for evaluation.

### 1.5. Measurement of plasma biochemistry

Plasma ALT levels were measured by FUJI DRI-CHEM 7000 (Fujifilm Corporation).

### 1.6. Measurement of liver biochemistry

#### 1.6.1. Measurement of liver lipid content

Liver total lipid extracts were obtained by Folch’s method [11]. Liver samples were homogenized in chloroform-methanol (2:1, v/v) and incubated overnight at room temperature. After washing with chloroform-methanol-water (8:4:3, v/v/v), the extracts were evaporated to dryness and dissolved in isopropanol. Liver triglyceride content was measured by the Triglyceride E-test (Wako Pure Chemical Industries, Ltd., Japan). Liver free fatty acid content was measured by the NEFA C-test (FUJIFILM Wako Pure Chemical Corporation).

### 1.7. Histological analysis

Sections (4 μm) were cut from paraffin blocks of liver tissue using a rotary microtome (Leica Microsystems). After sectioning, each slide was coded with a number for blinded evaluation by the pathologist. Each number was generated using the RAND function of Excel software, sorted in ascending order and assigned to slides.

#### Histological analyses

For Hematoxylin and Eosin (HE) staining, sections were cut from paraffin blocks of liver tissue prefixed in Bouin’s solution and stained with Lillie-Mayer’s Hematoxylin (Muto Pure Chemicals Co., Ltd., Japan) and eosin solution (Wako Pure Chemical Industries).

The NAFLD Activity Score (NAS) was calculated according to the criteria of Kleiner [10], as shown in Table 1. For NAS, bright field images of HE-stained sections were captured using a digital camera (DFC295; Leica, Germany) at 50- and 200-fold magnifications. Steatosis score in 1 section/mouse (representative 1 field at 50-fold magnification), inflammation score in 1 section/mouse (representative 1 field around the central vein at 200-fold magnification) and ballooning score in 1 section/mouse (representative 1 field around the central vein at 200-fold magnification) were estimated.

**Table 1:**
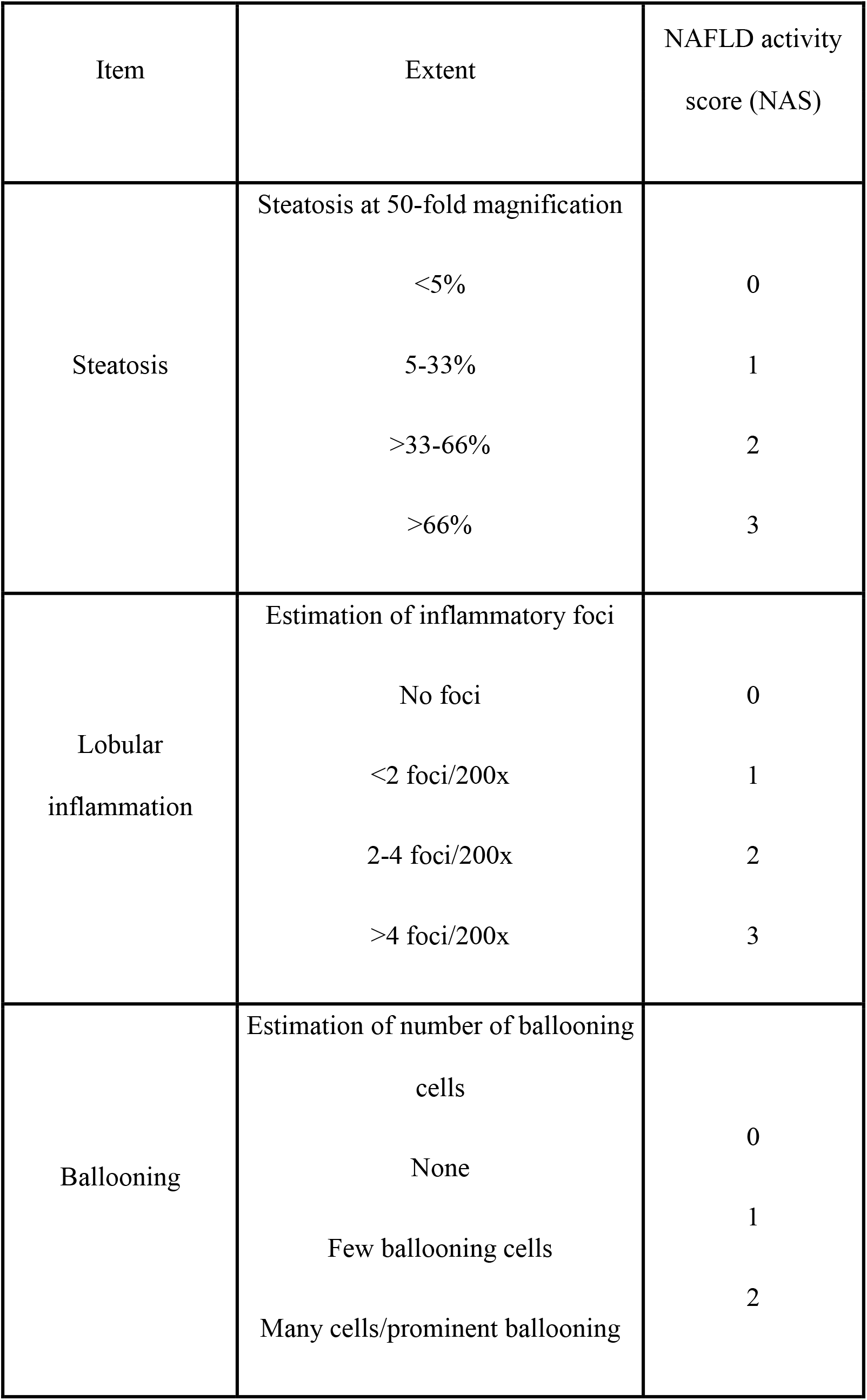
Definition of NAFLD Activity Score (NAS) components

To visualize collagen deposition, Bouin’s fixed liver sections were stained using picro-Sirius red solution (Waldeck, Germany). Briefly, sections were deparaffinized and hydrophilized with xylene, 100-70% alcohol series and RO water, and then treated with 0.03% picro-Sirius red solution (Cat No.: 1A-280) for 60 minutes. After washing with 0.5% acetic acid solution and RO water, stained sections were dehydrated and cleared with 70-100% alcohol series and xylene, then sealed with Entellan^®^ new (Merck, Germany) and used for observation.

For immunohistochemistry, sections were cut from frozen liver tissues embedded in Tissue-Tek OCT compound and fixed in acetone. Endogenous peroxidase activity was blocked using 0.03% H_2_O_2_ for 5 minutes, followed by incubation with Block Ace (Dainippon Sumitomo Pharma Co. Ltd., Japan) for 10 minutes. The sections were incubated with anti-F4/80 antibody at 4°C overnight. After incubation with a secondary antibody, enzyme-substrate reactions were performed using 3, 3’-diaminobenzidine/H_2_O_2_ solution (Nichirei Bioscience Inc., Japan). The primary antibody used was monoclonal antibody to mouse macrophages (BMA Biomedicals) at a dilution of 100 folds. The peroxidase-based detection system, VECTASTAIN ABC KIT (Vector Laboratories) was used as the secondary antibody staining system.

For quantitative analysis of the fibrosis area and inflammation area, bright field images of Sirius red-stained and F4/80-immunostained sections were captured around the central vein using a digital camera (DFC295; Leica, Germany) at 200-fold magnification, and the positive areas in 5 fields/section were measured using ImageJ software (National Institute of Health, USA).

### 1.8. Statistical Analysis

Statistical analyses were performed using Prism Software 6 (GraphPad Software, USA). Comparisons were made between the following groups using the Bonferroni multiple comparison test:

1. Group 1 (Vehicle) vs. Group 2 (AFO-202 Beta Glucan), Group 3 (N-163 Beta Glucan), Group 4 (AFO-202 Beta Glucan+ N-163 Beta Glucan) and Group 5 (Telmisartan) *P* values < 0.05 were considered statistically significant. Results were expressed as mean ± SD.

A trend or tendency was assumed when a one-sided t-test returned *P* values < 0.1. Comparisons were made between the following groups:

1. Group 1 (Vehicle) vs. Group 2 (AFO-202 Beta Glucan)
2. Group 1 (Vehicle) vs. Group 3 (N-163 Beta Glucan)
3. Group 1 (Vehicle) vs. Group 4 (AFO-202 Beta Glucan+ N-163 Beta Glucan)
4. Group 1 (Vehicle) vs. Group 5 (Telmisartan)

## Results

There was no significant difference in the body weight and liver weight between the groups (Figure 1). The mean ± SD of body weight was 20.4 ± 1.9 g in Group 1, 20.3 ± 1.3 g in Group 2, 20.2 ± 1.5 g in Group 3, 19.6 ± 2.1 g in Group 4 and 17.8 ± 0.9 g in Group 5. The mean ± SD liver weight was 1552 ± 162 mg in Group 1, 1552 ± 92 mg in Group 2, 1565 ± 182 mg in Group 3, 1474 ± 197 mg in Group 4 and 1181 ± 123 mg in Group 5.

**Figure 1:**
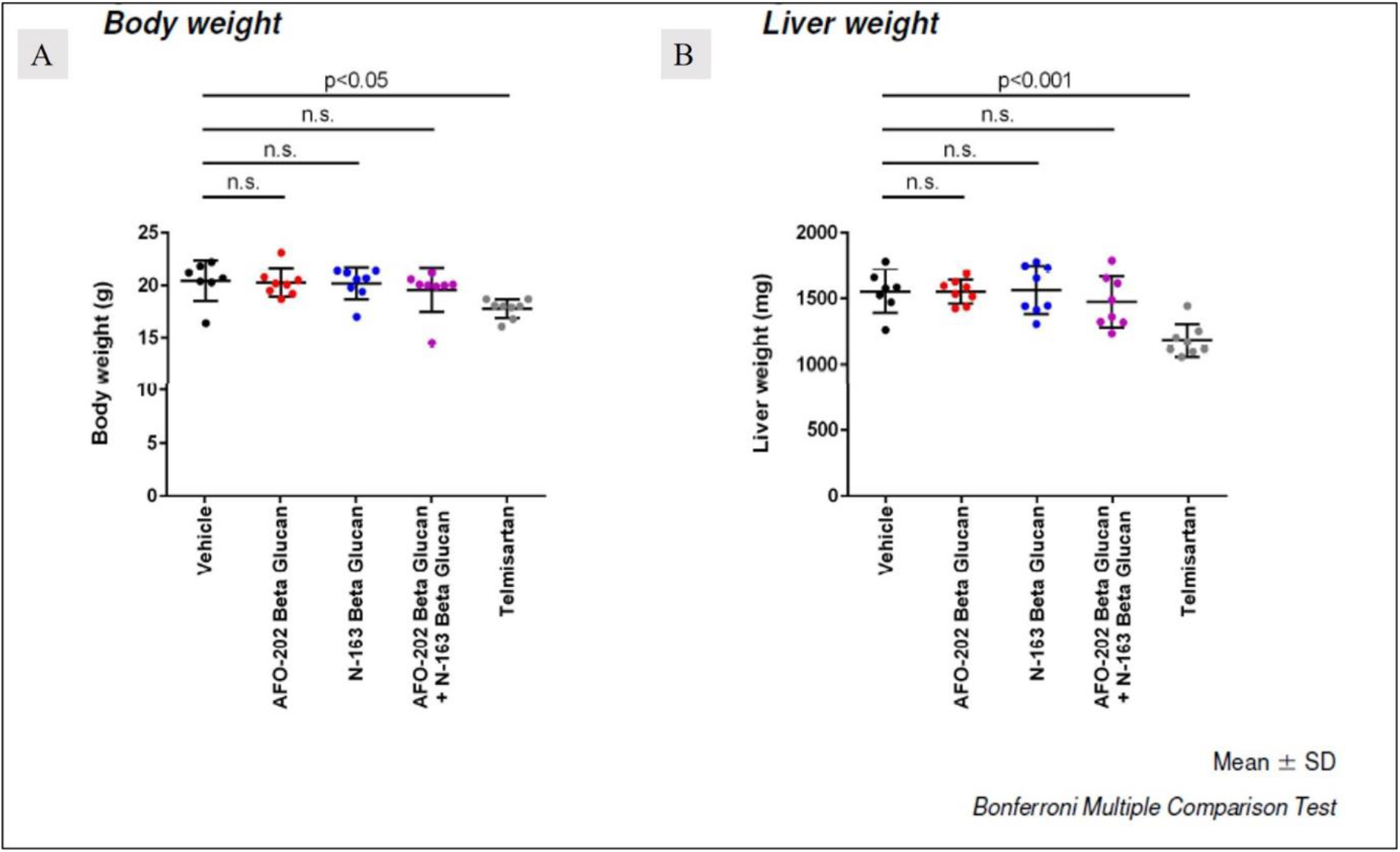
Body weight and liver weight showing no significant difference between the groups; telmisartan brings down total body weight and liver weight compared to other groups

Plasma ALT levels were lowest in the telmisartan group (Mean ± SD = 36 ± 7 U/L), followed by Group 3 (N-163) (Mean ± SD = 44 ± 8 U/L) (Figure 2).

**Figure 2:**
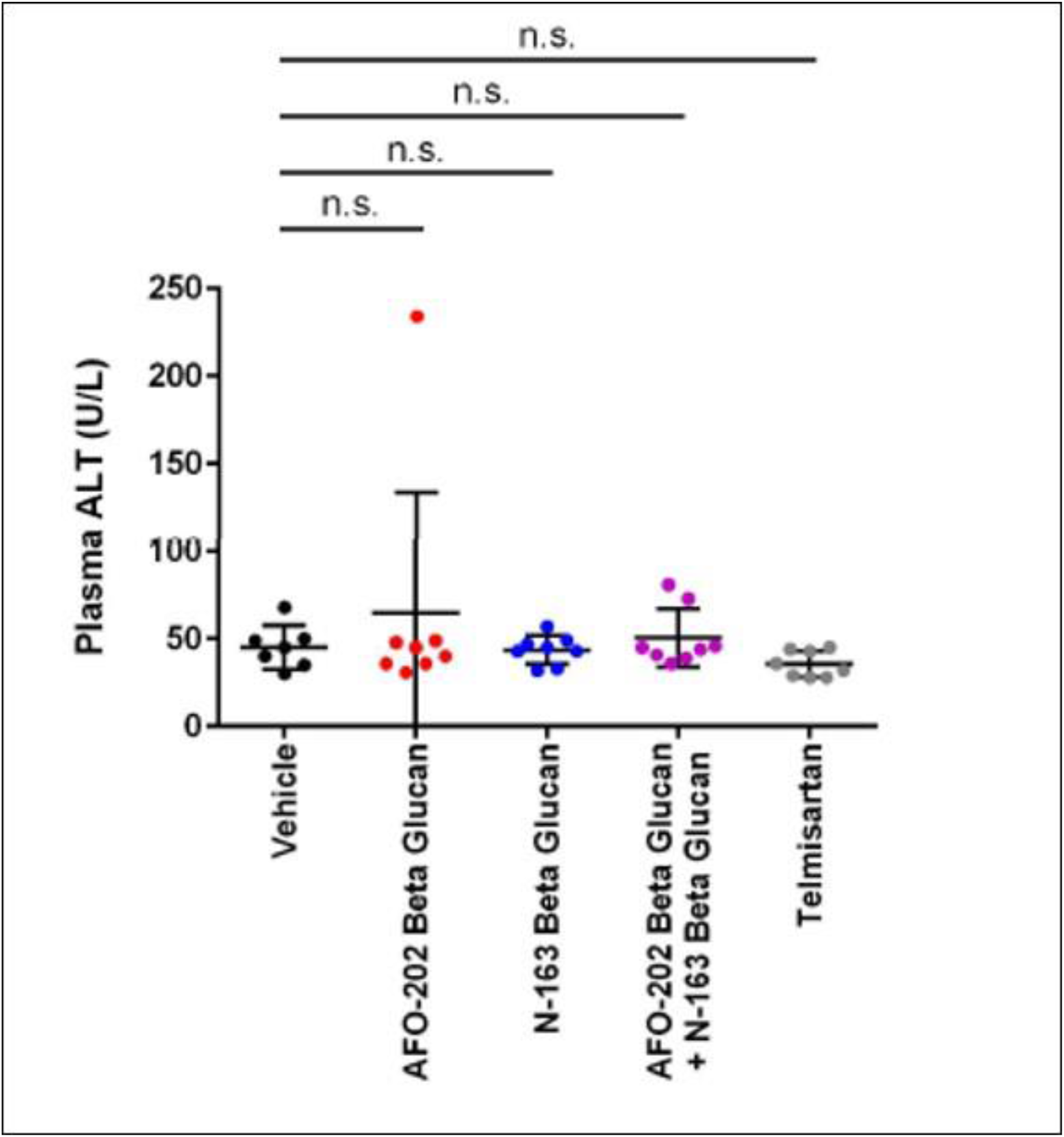
Plasma ALT (mg/dL) levels were decreased in the telmisartan and N-163 groups compared to the other groups

Representative photomicrographs of Sirius red-stained liver sections are shown in Figure 3. Liver sections from the vehicle group showed increased collagen deposition in the pericentral region of liver lobule. Sirius red-stained images to assess liver damage showed significantly decreased positive staining area in the AFO-202+N-163 and N-163 groups (p<0.05) compared with all the other groups (average positive stained area, AFO-202-0.80 ± 0.22 AFO-202+N-163: 0.65 ± 0.25; N-163: 0.56 ± 0.12; telmisartan: 0.59 ± 0.20; and vehicle: 0.96 ± 0.22) (Figure 3, 4).

**Figure 3.**
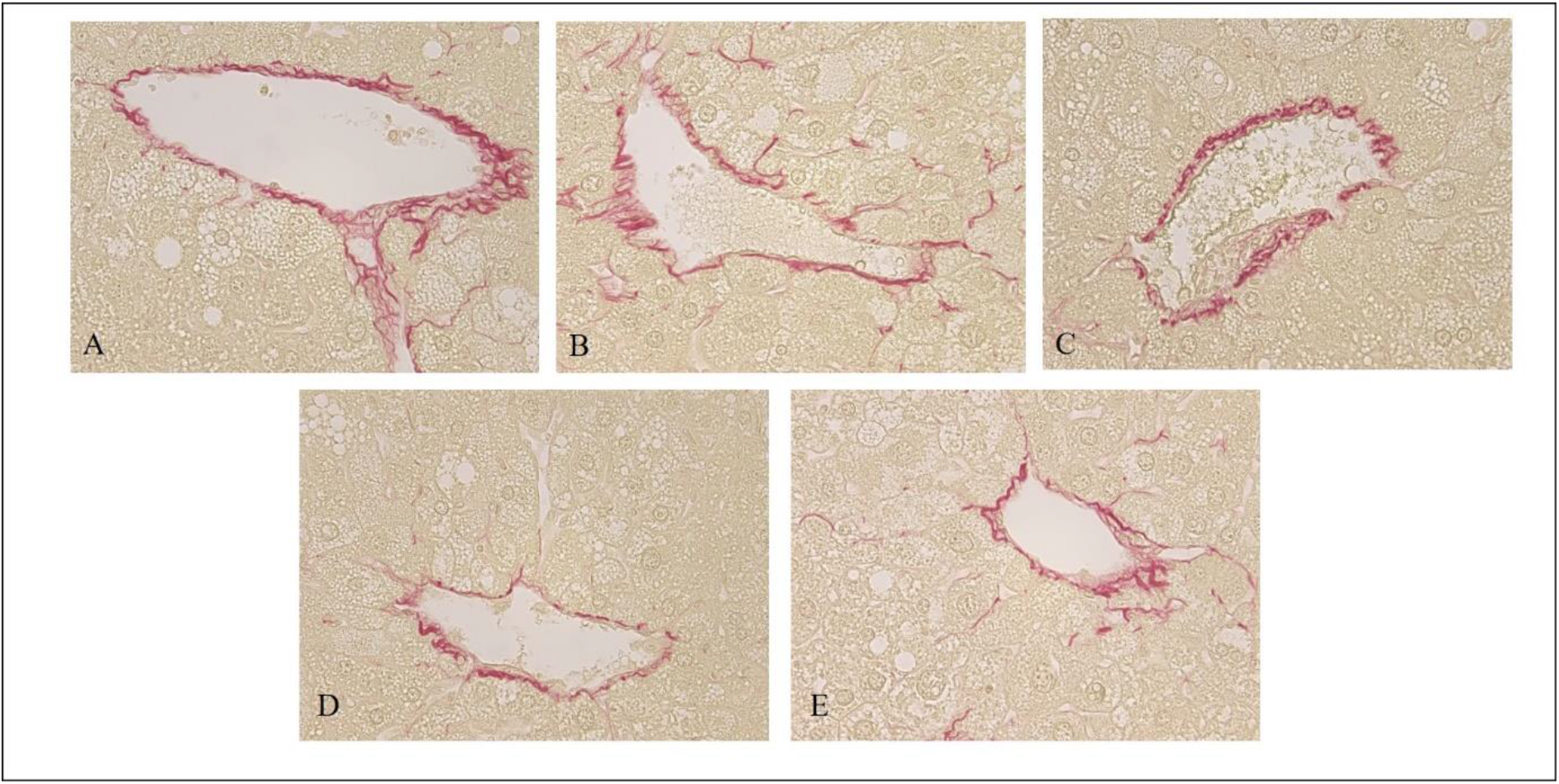
Hepatic Fibrosis evaluated with Sirius red staining of A. Vehicle; B. AFO-202; C. N-163; D. AFO 202+N-163 and E. Telmisartan with AFO-202+N-163 (D) and N-163 (C) showing significantly decreased positive staining area compared with all the other groups (Magnification: x400)

**Figure 4:**
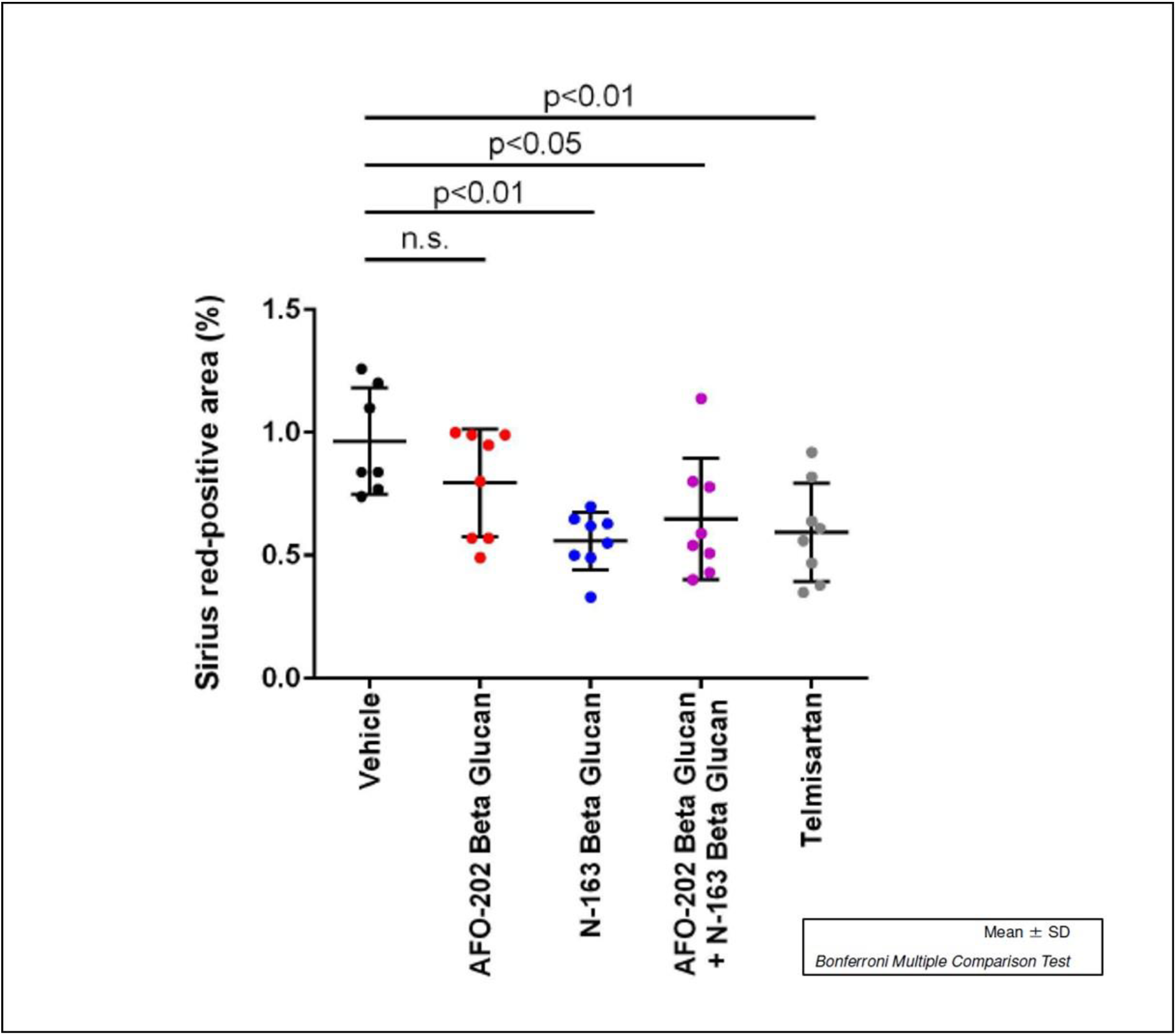
Average positive stained area for fibrosis showing significantly decreased positive staining area in the AFO-202+N-163 and N-163 groups compared with all the other groups

In H and E staining, liver sections from the vehicle group exhibited micro- and macro vesicular fat deposition, hepatocellular ballooning and inflammatory cell infiltration. All the beta glucan treatment groups showed significant decreases in NAS compared with the Vehicle group (Figure 5).

**Figure 5:**
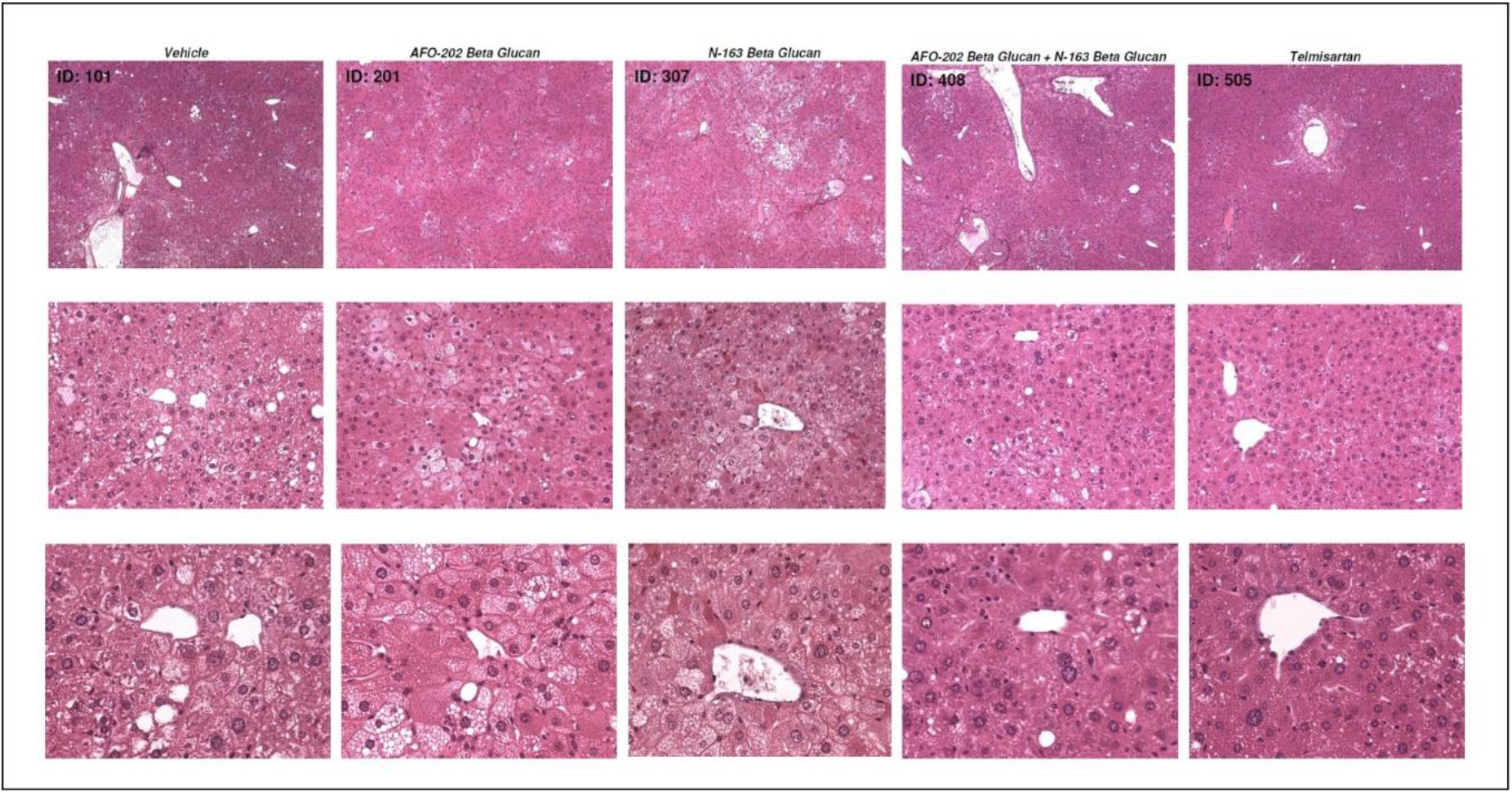
Representative photomicrographs of HE-stained liver sections: Upper panel: Magnification = x50; Middle panel: Magnification = x200; Lower panel: Magnification = x400

The telmisartan, N-163 and AFO-202 +N-163 groups of mice had a significantly lower NAFLD activity score (NAS) compared with untreated and vehicle treated groups of mice (Mean score, telmisartan: 2.6 ± 0.7; AFO-202+N-163: 3.3 ± 1.0; N-163: 3.5 ± 0.5; AFO-202: 3.3 ± 0.9 and vehicle: 4.6 ± 0.5). The inflammation score was significantly decreased in the AFO-202+N-163 and N-163 groups compared with the telmisartan group (Figures 5,6). Ballooning and steatosis score was decreased most in the telmisartan group, but a decrease in ballooning and steatosis compared with vehicle treated mice was observed in the AFO-202 beta glucan groups (Figure 5, 6).

**Figure 6:**
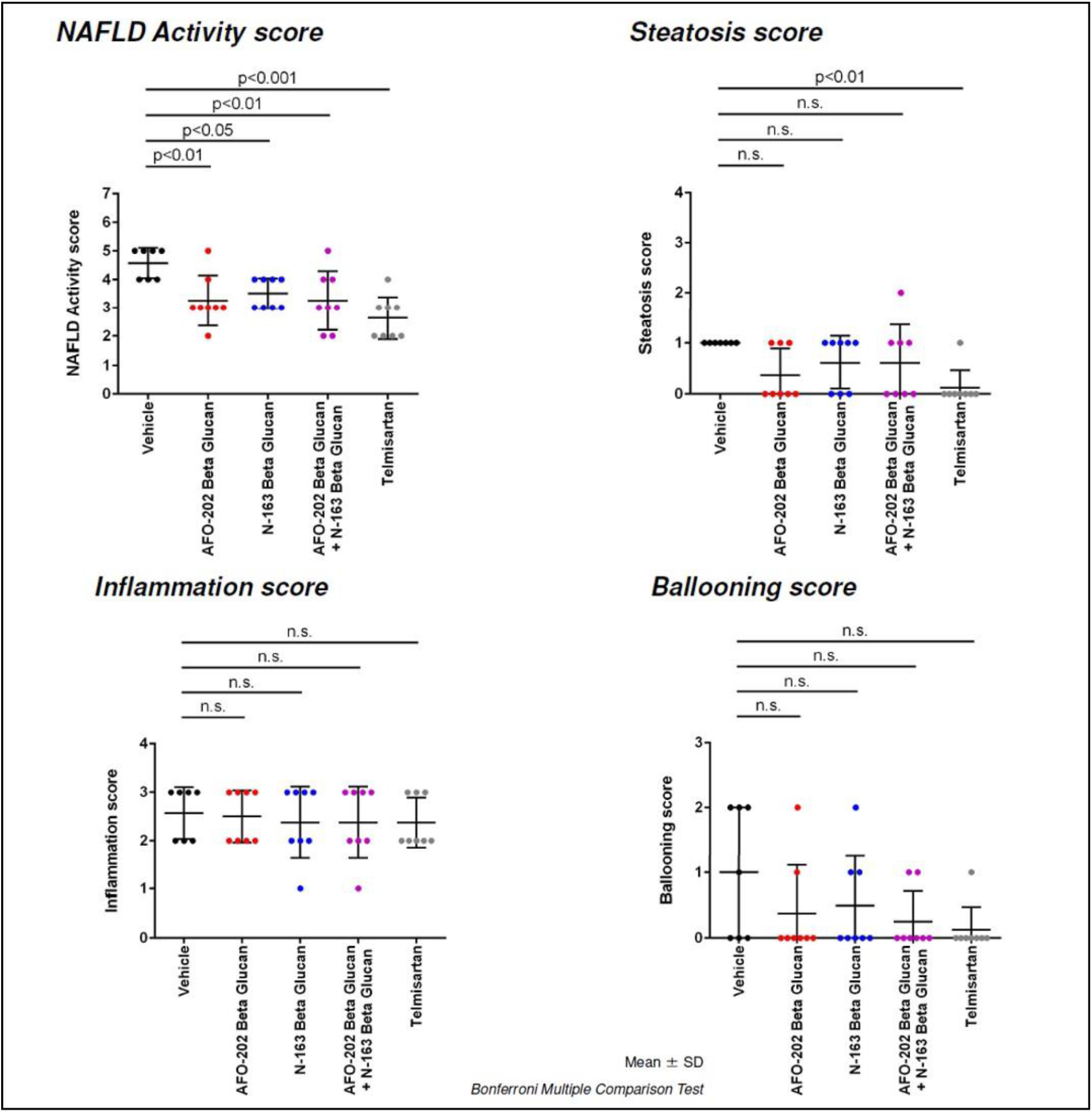
NAFLD activity score (NAS), steatosis, inflammation and ballooning scores in the various groups based on H and E staining

Representative photomicrographs of F4/80-immunostained liver sections are shown in Figure 7. F4/80 immunostaining of liver sections from the vehicle group demonstrated accumulation of F4/80+ cells (macrophages associated with inflammation) in the liver lobule. F4/80 immunostaining score for these inflammatory macrophages was least in the N-163 group compared to the other groups (Figure 7, 8)

**Figure 7:**
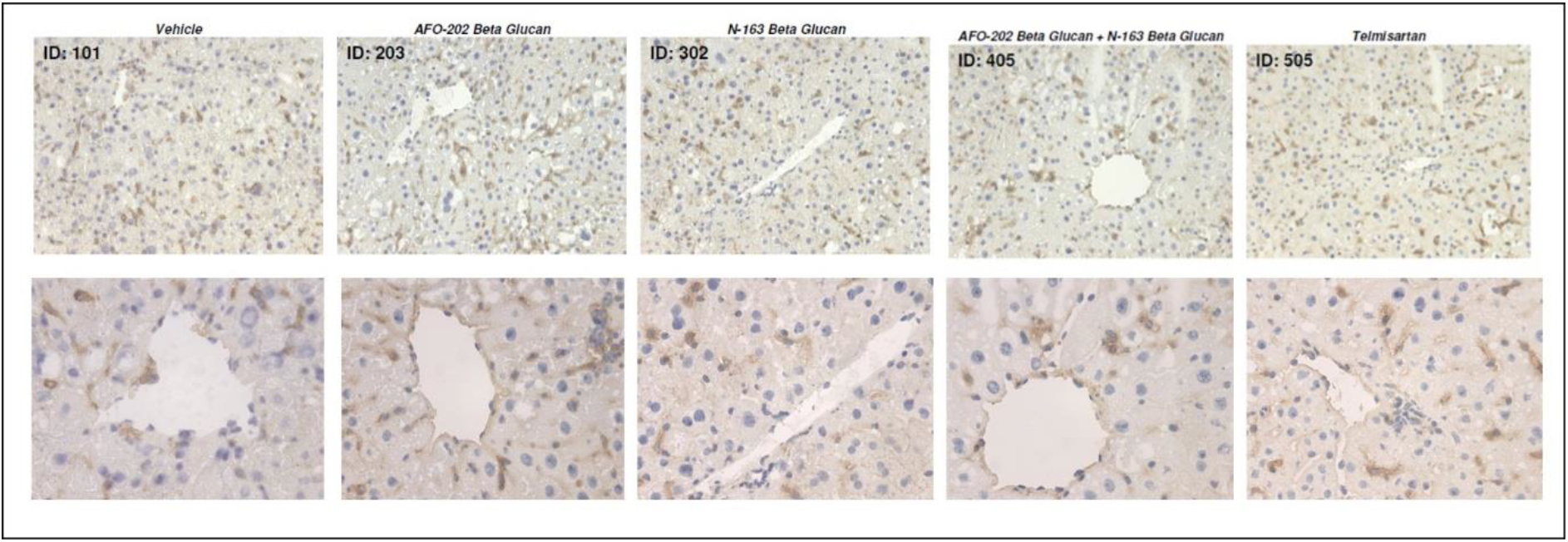
A. Representative photomicrographs of F4/80-immunostained liver sections Upper panel: Magnification: x200; Lower panel: Magnification: x400

**Figure 8:**
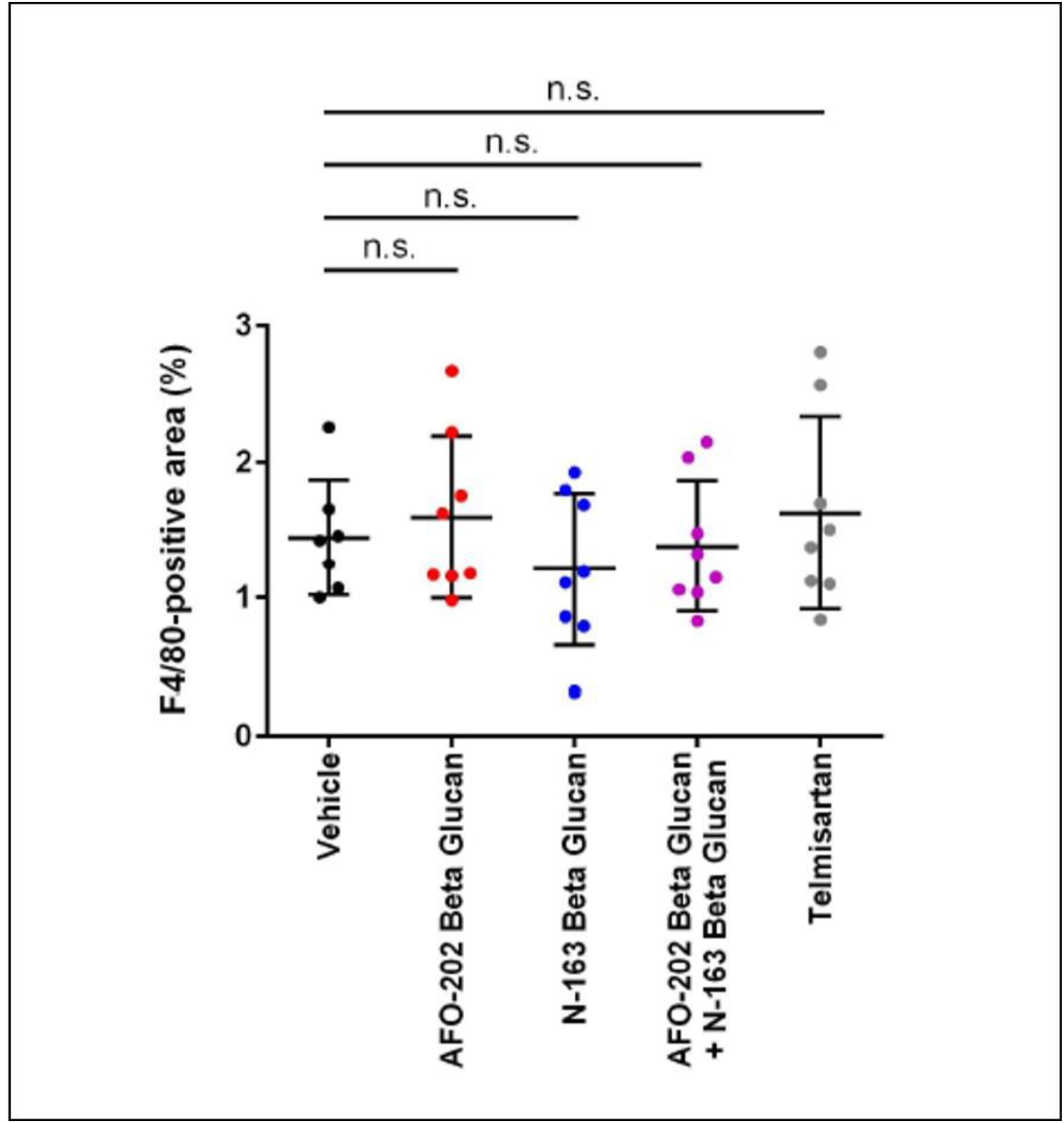
Scoring of the inflammation area by F4/80 immunostaining which shows that the macrophages associated inflammation was least in the N-163 group compared to the other groups

## Discussion

NASH or NAFLD is a serious chronic liver disease that at first is a metabolic imbalance leading to accumulation of fat in the liver, and the inflammatory response to excess fat accumulation leads gradually to fibrosis, jeopardizing the liver function, and beyond this, if the chronic inflammation continues, it may lead to cirrhosis and hepatocellular carcinoma [12]. Increased plasma glucose and lipid levels contribute to direct lipid deposition in the liver and lead to systemic inflammation which contributes to the development and worsening of NAFLD [13]. A strategic approach to NAFLD therefore would be first to address the metabolic imbalance, which, based on our earlier findings, could be managed by administration of AFO-202 beta glucan [7-9], while the resolution of the already established fibrosis could be addressed by N-163 beta glucan, as shown in the present study. The present study has also proven that the combination of AFO-202 and N-163 is effective to address the chronic-inflammation-fibrosis cascade, preventing the culmination in cirrhosis or progress to carcinoma.

In this study, the effects of AFO-202 and N-163 Beta 1,3-1,6 glucans were tested individually and in combination in the STAM mice model of NASH. The decrease in body weight and liver weight was significantly reduced only in the telmisartan group (Figure 1). The inflammation and ballooning scores were decreased mainly in the AFO-202 beta glucan groups, indicating that beta glucans may act as an anti-inflammatory protective agent against NASH progression (Figure 5). AFO-202 beta glucan has been shown to decrease inflammation-related cytokines in previous studies [14]. This is further substantiated in the present study. However, fibrosis, which is the outcome of inflammation (Figure 4) and macrophage associated inflammation (F4/80 immunostaining) was reduced mainly in the N-163 group (Figure 8), and the steatosis and NAS were decreased in the AFO-202 +N-163 groups as effectively as in the telmisartan group (Figures 6), indicating their application as an anti-fibrotic treatment agent in NASH. Beta glucans have been reported to help in alleviating obesity by acting on modulating transcription factor peroxisome proliferator-activated receptor (PPAR)-γ [15]. This could be one probable mechanism behind the hepatoprotective anti-inflammatory and anti-fibrotic effects of AFO-202 and N-163 in the current study. Gut microbiota, which are dysregulated in metabolic syndrome, diabetes and dyslipidaemia, also lead to NASH by production of endotoxins. The prebiotic effects of the AFO-202 and N-163 beta glucans could also contribute to NASH alleviation by helping with gut microbiota’s beneficial alteration [1], which needs further validation.

AFO-202, N-163 beta glucans and their combination are known food supplements with established safety after decades of human consumption [14] in contrast to a pharmacological agent such as telmisartan adds to their potential for managing NAFLD. The other beneficial effects of these beta glucans on obesity, diabetes and dyslipidaemia [7-10, 15,16] also provides a strong rationale for their therapeutic use in NASH related diseases.

Furthermore, the mechanisms of the minute specific details though may be difficult, so additional evaluation of (i) gene expression for fibrotic and inflammatory markers in the STAM model after AFO-202, N-163 beta glucans administration could shed light on intricacies for a better understanding of these beta glucans in NASH, while (ii) evaluating common markers of tissue and organ fibrosis to other organ diseases such as PPAR-γ TGFb, TNFα, MCP-1, α-SMA, TIMP-1 [17, 18] could add value to examining the possibilities of an extended application of these BRMGs in lung and kidney fibrosis as well. IL-6, having been already shown to be decreased by the AFO-202 beta glucan, which is a key cytokine implicated in inflammatory and fibrosis mechanisms [19] of lung, liver and kidney [20], is a specific biomarker worth evaluation in further studies.

Having been proven to be safe for human consumption as a food supplement, these two novel beta glucans, AFO-202 studied for 25 years and N-163, a larger multicentre study in NASH/NAFLD patients would be appropriate. However, one has to keep in mind the limitations of this study, including the fact that though this STAM model recapitulated human fatty liver disease to a great extent, there are still differences in the immunological mechanisms mediating inflammation between humans and mice, with human neutrophil-attracting chemokine IL-8 having no direct analogue in mice and differences in the corresponding immune cell subsets between mice and humans [21].

## Conclusion

This study was a comprehensive preclinical evaluation demonstrating the hepatoprotective anti-fibrotic effects of N-163, anti-inflammatory effects of AFO-202 beta glucan and a combination of these two biological response modifier glucans in decreasing the NAS score in an established NASH model of fatty liver disease, STAM. Considering the safety of these two food supplements, a larger clinical study in NASH patients is recommended, and further research on these beta glucans and their beneficial effects through gene expression and common biomarkers of tissue and organ fibrosis is worthwhile, as the fundamental mechanisms of fibrosis in other organs such as the kidney and lungs have common mechanisms.

## Acknowledgements

The authors thank

a. Mr. Yasunori Ikeue, Mr. Mitsuru Nagataki and Mr. Takashi Onaka, (Sophy Inc, Kochi, Japan), for necessary technical clarifications.
b. Mr. Yoshio Morozumi, Ms. Yoshiko Amikura of GN Corporation, Japan for their liaison assistance with the conduct of the study.
c. Loyola-ICAM College of Engineering and Technology (LICET) for their support to our research work.

